# Comparison of everting sutures and the lateral tarsal strip with or without everting sutures for involutional lower eyelid entropion: A meta-analysis

**DOI:** 10.1101/2020.12.30.424787

**Authors:** Gyudeok Hwang, Hyo Sin Kim, Jiyoung Lee, Ji-Sun Paik

## Abstract

There are three pathophysiologies of involutional entropion, vertical laxity (VL), horizontal laxity (HL), and overriding of the preseptal orbicularis. The effects of methods to correct VL only, HL only, or both VL and HL in patients with involutional entropion were compared using the published results of randomized controlled trials (RCTs).

To find RCT studies that investigated methods to correct involutional entropion, a systematic search was performed from database inception to April 2020 in the Medline, EMBASE, and Cochrane databases. Two independent researchers conducted the literature selection and data extraction. Evaluation of the quality of the reports was performed using the Cochrane Collaboration tool for assessing the risk of bias (ROB 2.0). The data analysis was conducted according to the PRISMA guidelines using Review Manager 5.3.

Two RCT studies were included in this meta-analysis. Surgery for involutional entropion was performed on a total of 109 eyes. Everting sutures (ES) were used on 57 eyes and lateral tarsal strips (LTS) or combined procedures (LTS + ES) were performed on 52 eyes. At the end of the follow-up periods, involutional entropion recurred in 18 eyes (31.6%) in the ES group and three eyes (5.8%) in the LTS +/- ES group. Analysis of the risk ratio showed that the LTS +/- ES method significantly lowered the recurrence rate compared to using ES only (P = 0.007).

Performing LTS +/- ES effectively lowered the recurrence rate of involutional entropion compared to ES alone. However, some patients cannot tolerate more invasive corrections such as LTS. Therefore, sequential procedures, in which ES is performed first and then when entropion recurs LTS +/- ES is performed, or another methods depending upon the degree of HL may be used.

## Introduction

Entropion is an eyelid malposition where the eyelid margin and eyelashes are turned toward the eyeball. Entropion is divided into four types, cicatricial, congenital, acute spastic, and involutional [1]. Involutional entropion, also known as senile entropion, is most commonly observed in general ophthalmic practice and increases in incidence with age [2, 3]. Also, the incidence of entropion in Asians is higher than in non-Asians [4]. Patients with involutional entropion complain of dry eye syndrome, superficial punctate keratopathy, chronic blepharitis, and chronic conjunctivitis [2]. Non-surgical therapies such as the use of lubricating ointment, eyelid taping, and botulinum toxin injections are used for the treatment of involutional entropion, but most are temporary treatments while the patient awaits eyelid surgery, which is the definitive treatment [1, 5]. The causative factors of involutional entropion are 1) vertical laxity of the lower eyelid, 2) horizontal laxity of the lower eyelid, and 3) overriding of the preseptal orbicularis oculi muscle (OOM) [6, 7].

Various surgical methods have been attempted to correct each causative factor of involutional entropion. The methods to correct vertical laxity of the lower eyelid include everting sutures (ES), the Quickert procedure, the Weis procedure, the Jones procedure, the Hotz procedure, lower eyelid retractor advancement, and Bick’s procedure. Lateral tarsal strips (LTS) and lateral wedge resection are used to correct horizontal laxity of the lower eyelid and OOM transposition is a method of correcting overriding of the pre-septal OOM. In addition, procedures combining these methods, such as ES with LTS or lower eyelid retractor advancement with LTS, are also performed [5]. We performed a meta-analysis of randomized controlled trial (RCT) results of the recurrence and complication rates after procedures conducted to correct vertical laxity only, horizontal laxity only, or both in patients with involutional entropion. In addition, we summarized all RCTs performed for involutional entropion and the results.

## Materials and Methods

### Search strategy and study selection

To identify RCTs comparing ES and LTS with ES or LTS only (LTS+/-ES) for involutional entropion, a systematic search was performed from database inception to April 2020 in the Medline, EMBASE, and Cochrane databases. To briefly describe the search strategy, the target diseases ‘involutional lower eyelid entropion’, ‘involutional entropion’, and ‘senile lower eyelid entropion’, ‘senile entropion’, which are used interchangeably, were used as the search terms. Among the searched articles, all RCTs targeting involutional lower eyelid entropion were found and a meta-analysis was performed only for RCTs comparing ES and LTS +/- ES. The selection criteria for the relevant studies were (1) randomized controlled trials, (2) patients with involutional lower eyelid entropion, and (3) patients treated with ES vs. LTS +/- ES. There was no restriction on the publication language of the articles if the abstracts were in English. The exclusion criteria were (1) non-RCT trials, (2) other diseases, and (3) interventions other than ES vs. LTS +/- ES.

### Data extraction

Excluding data duplication, 351 articles were searched. Two investigators reviewed the titles and abstracts independently and selected 158 potentially eligible studies. After that, a full-text review was conducted. Disagreements in the literature selection were resolved by discussion between the two investigators. Finally, two articles were selected [8, 9] (Fig 1).

**Fig 1.**
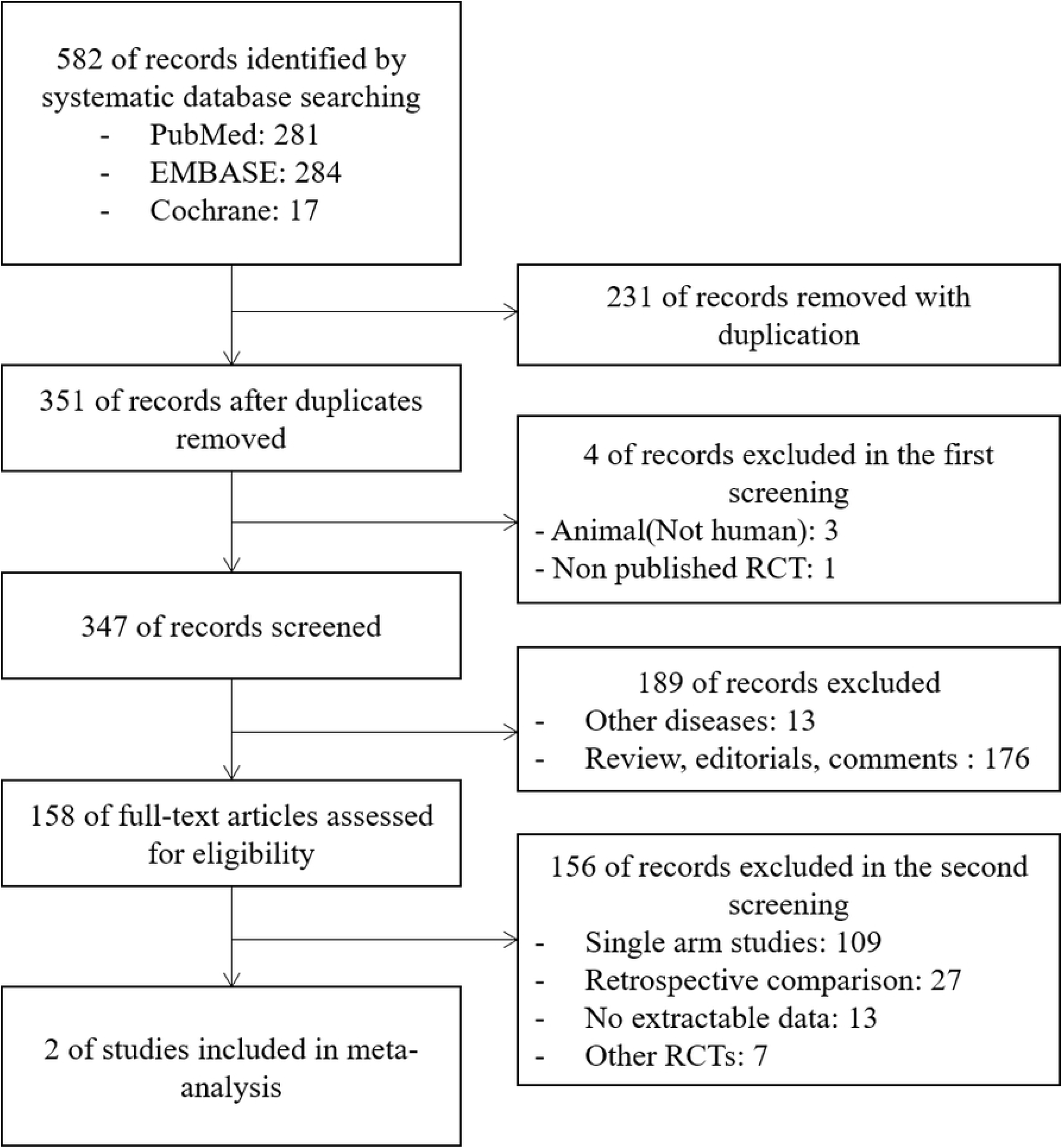
Flow diagram of studies included in this review.

Information on the country where the study was conducted, the number and characteristics of the participants, the follow-up period, and the number of surgeons was extracted from each article. The outcomes were the number of recurrent patients and complications at each follow-up point. In this paper, the term “end of study follow-up point” refers to the end of the trials or the last follow-up point of each trial.

### Quality assessment

A quality assessment of the included RCTs was performed with the Cochrane Collaboration tool for assessing the risk of bias (ROB; ROB 2.0, version of 22 August 2019). Two investigators assessed the ROB independently and resolved divergences through a consensus. Although blinding was not mentioned in the abstract and full text and it is thought that the participants in the RCT studies knew the aligned interventions in advance, the domain of ‘deviations from the intended interventions’ was evaluated as ‘some concerns’ if the analysis process seemed to be performed as the protocol. If the follow-up loss was more than 10% and analysis of the participant characteristics was not presented, the domain of “missing outcome data” was evaluated as “some concerns.” Although it was not described in the abstracts and full texts, the domain of “measurement of the outcome” was evaluated as high risk if it was thought that the assessors knew the participants’ aligned interventions in advance. The overall ROB for each trial is presented in Table 1. The ROB for each domain analyzed using Review Manager 5.3 is presented in the graph in Fig 2 and the ROB summary is shown in Fig 3.

**Table 1.**
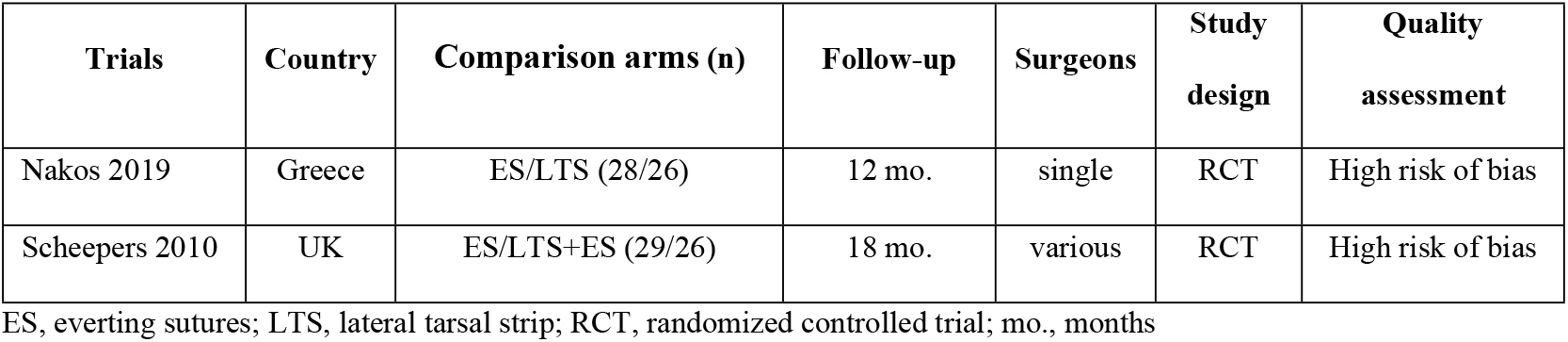
Characteristics of the included trials.

| **Trials** | **Country** | **Comparison arms (n)** | **Follow-up** | **Surgeons** | **Study design** | **Quality assessment** |
| --- | --- | --- | --- | --- | --- | --- |
| Nakos 2019 | Greece | ES/LTS (28/26) | 12 mo. | single | RCT | High risk of bias |
| Scheepers 2010 | UK | ES/LTS+ES (29/26) | 18 mo. | various | RCT | High risk of bias |

ES, everting sutures; LTS, lateral tarsal strip; RCT, randomized controlled trial; mo., months

**Fig 2.**
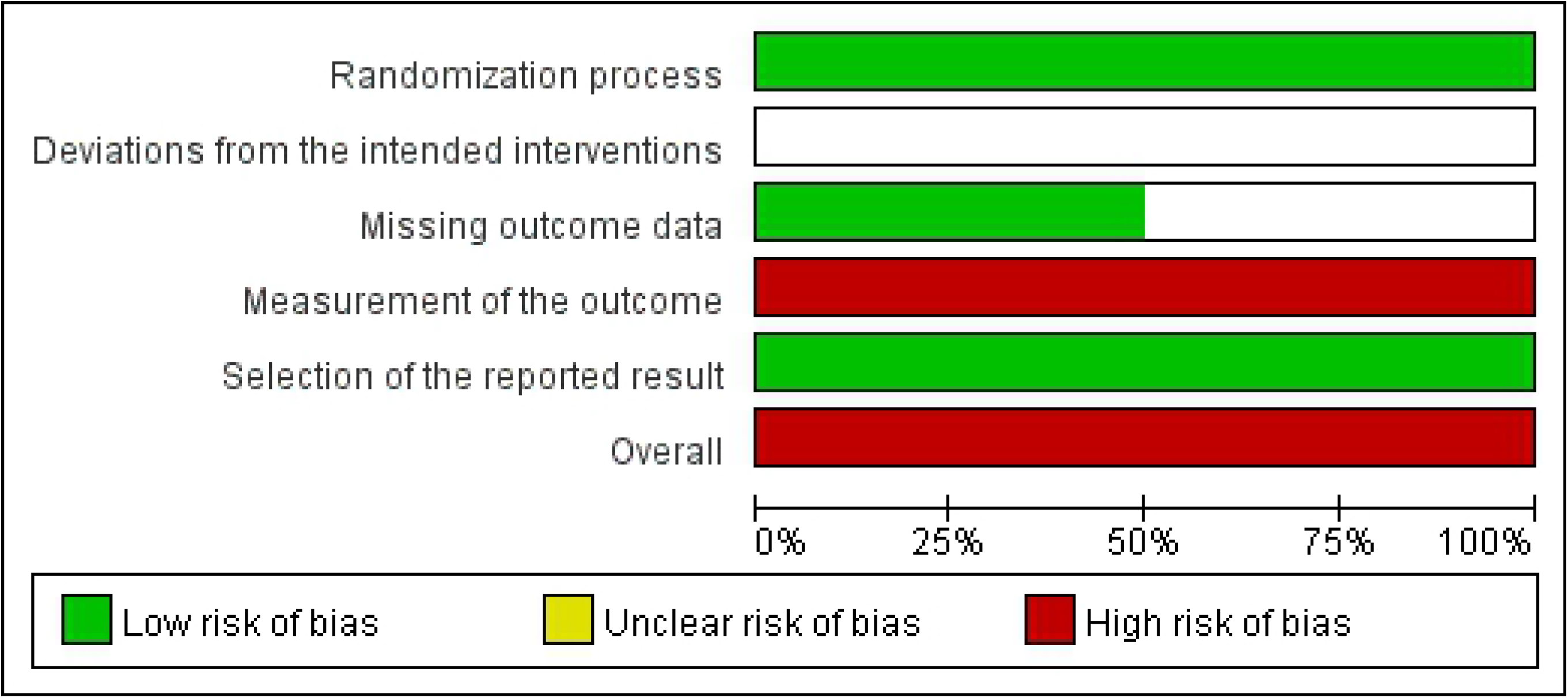
Risk of bias graph. The reviewing authors’ judgments on each risk of bias item in all included studies are presented as percentages. “Unclear risk of bias” means “some concerns.”

**Fig 3.**
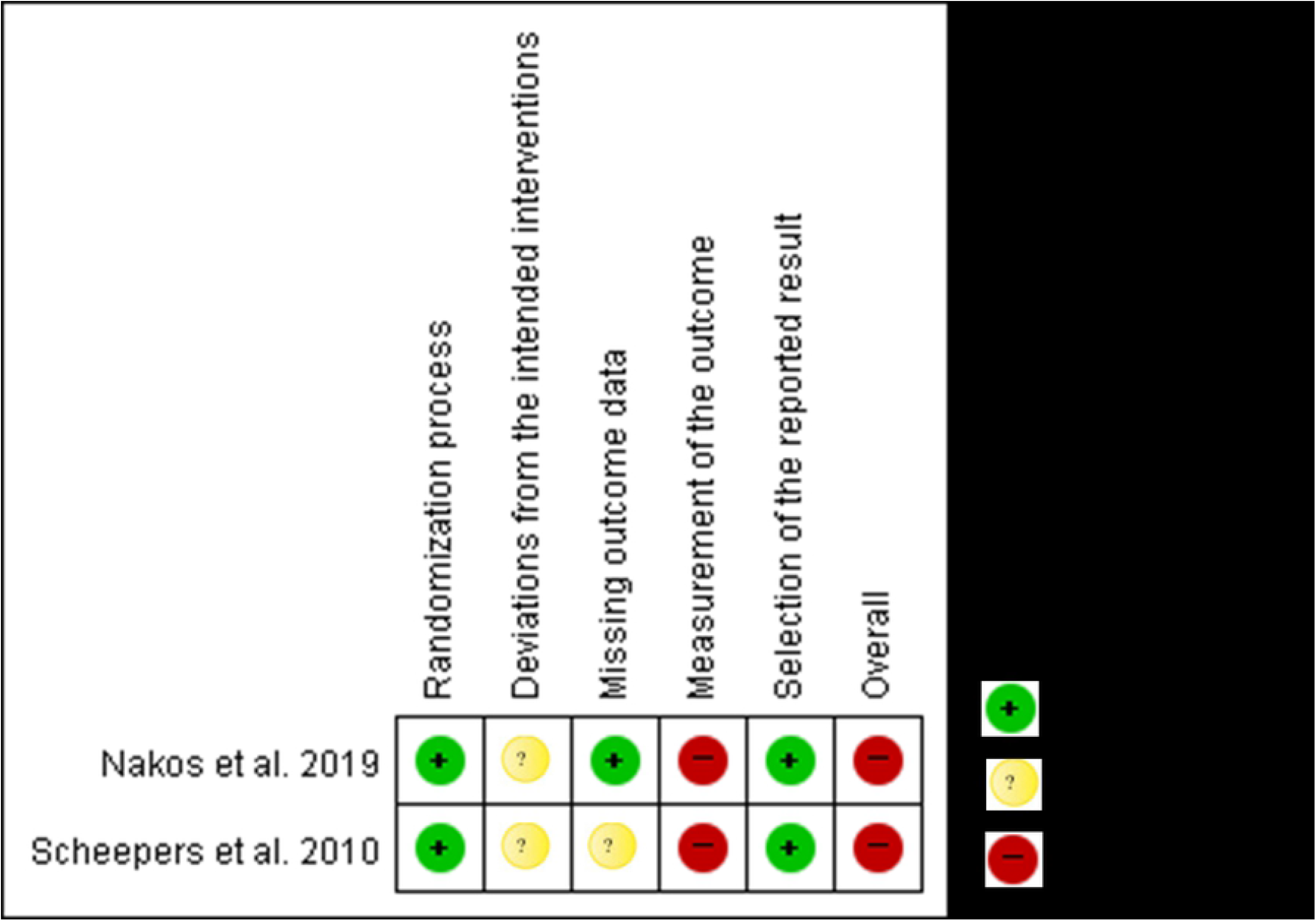
Risk of bias summary. The reviewing authors’ judgments on each risk of bias item in each included study.

### Statistical analyses

The outcome data were analyzed for recurrence and complication rates using Review Manager 5.3. In this meta-analysis, the random-effects model and the Mantel-Haenszel method were used because heterogeneity was suspected. The surgical methods used in each trial were not the same. One was ES vs. LTS and the other was ES vs. LTS + ES. The end of the study follow-up times were 12 months and 18 months. For these reasons, the assumption that the different studies were estimating different, yet related, intervention effects was applied [10]. Since the outcomes to be compared were dichotomous variables classified as the presence or absence of recurrence, a risk ratio (relative risk) was used for comparison. The I^2^ statistic was used to evaluate the degree of heterogeneity. Moderate heterogeneity was defined as an I^2^ > 25% and severe heterogeneity was indicated by an I^2^ > 75%.

## Results

### Characteristics of the included trials

There were two trials in the two selected articles. One trial compared ES vs. LTS + ES and the other one compared ES vs. LTS for involutional entropion (Table 1). Both trials were conducted in Europe, Greece, and the UK. Interventions were performed on a total of 109 eyes. To correct vertical laxity only, ES were used on 57 eyes. LTS used to correct horizontal laxity only or combined procedures to correct both horizontal laxity and vertical laxity with LTS and ES(LTS +/- ES) was performed on 52 eyes. The smallest sample size was 26 eyes, the largest sample size was 29 eyes, and the median sample size was 27 eyes. The follow-up periods in the trials were 12 months and 18 months.

None of the follow-up points in the two trials coincided with each other (Tables 2 and 3).

**Table 2.** Recurrences at each follow-up point.

| Trials | ES |  |  | LTS +/- ES |  |  |
| --- | --- | --- | --- | --- | --- | --- |
|  | n (%) |  |  | n (%) |  |  |
|  | 6 mo. | 12 mo. | 18 mo. | 6 mo. | 12 mo. | 18 mo. |
| Nakos 2019 | 8 (28.6) | 12 (42.9) |  | 1 (3.8) | 3 (11.5) |  |
| Scheepers 2010 |  |  | 6 (20.7) |  |  | 0 (0) |

ES, everting sutures; LTS, lateral tarsal strip; mo., months

**Table 3.** Complications at each follow-up point.

| Trials | ES |  |  | LTS +/- ES |  |  |
| --- | --- | --- | --- | --- | --- | --- |
|  | n (%) |  |  | n (%) |  |  |
|  | 6 mo. | 12 mo. | 18 mo. | 6 mo. | 12 mo. | 18 mo. |
| Nakos 2019 | 0 (0) | 0 (0) |  |  | 1 (3.8) |  |
| Scheepers 2010 |  |  | 2 (6.9) |  |  | 0 (0) |

ES, everting sutures; LTS, lateral tarsal strip; mo., months

In the Nakos trial (2019) [8], entropion recurred in 12 of 28 eyes (42.9%) that underwent ES and three of 26 eyes (11.5%) that underwent LTS after 12 months. In the Scheepers trial (2010) [9], recurrence was observed in six of the 29 eyes (20.7%) that underwent ES, and recurrence was not observed in 26 eyes treated with LTS + ES (0%) after 18 months.

In the Nakos trial (2019) [8], no complications occurred among 28 eyes that underwent ES (0%), and complications occurred in one of 26 eyes that underwent LTS (3.8%) after 12 months. In the Scheepers trial (2010) [9], complications occurred in two of the 29 eyes that underwent ES (6.9%) and none of the 26 eyes that underwent LTS + ES (0%) after 18 months.

### Quality assessment

The assessment of RCT quality using the Cochrane Collaboration tool (version 2.0; August 22, 2019) for assessing ROB found that both trials had a high ROB (Table 1). In the last two selected articles, there was no mention of blinding of the participants, surgeons, and assessors, and it seems likely that they knew about the intervention assignments.

### Analysis

The recurrence rate was compared at the end of the study follow-up points of each trial using Review Manager 5.3 (Fig 4). At the end of the study follow-up points, 18 eyes (31.6%) in the ES group and three eyes (5.8%) in the LTS +/- ES group experienced recurrences. The risk ratio for recurrence between the ES group and the LTS +/- ES group was 4.37, and the recurrence rate in the LTS +/- ES group was significantly lower than that in the ES group (95% confidence interval: 1.51 to 12.64 P = 0.007). Recurrences in the two trials showed low heterogeneity with I^2^ = 0%.

**Fig 4.**
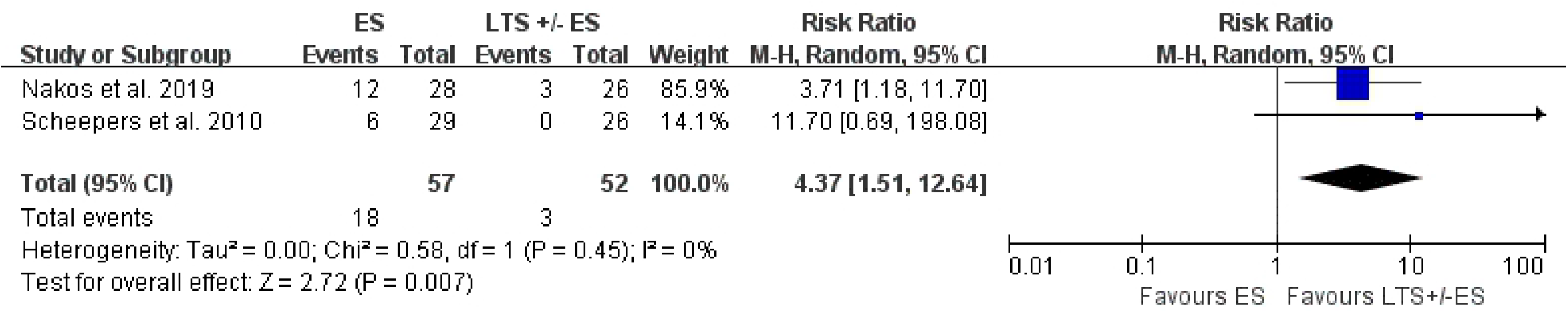
Forest plots of the effect sizes for recurrences at the end of the study follow-up points.

In the Nakos [8] and Scheepers trials [9], the risk ratios for recurrence between the ES group and LTS +/- ES group were 3.71 (95% confidence interval (CI): 1.18 to 11.70, P = 0.02) and 11.70 (95% CI: 0.69 to 198.08, P = 0.09), respectively.

The complications were compared at the end of the study follow-up points of each trial (Fig 5). At the end of the study follow-up points, complications occurred in two eyes in the ES group (3.5%) and one eye in the LTS +/- ES group (1.9%), respectively. The risk ratio for complications between the ES group and the LTS +/- ES group was 1.24. There were no statistically significant differences in the risk of complications between the two groups (95% confidence interval: 0.09 to 17.11 P = 0.87). Moderate heterogeneity was observed, with an I^2^ = 31% for complications in the two trials.

**Fig 5.**
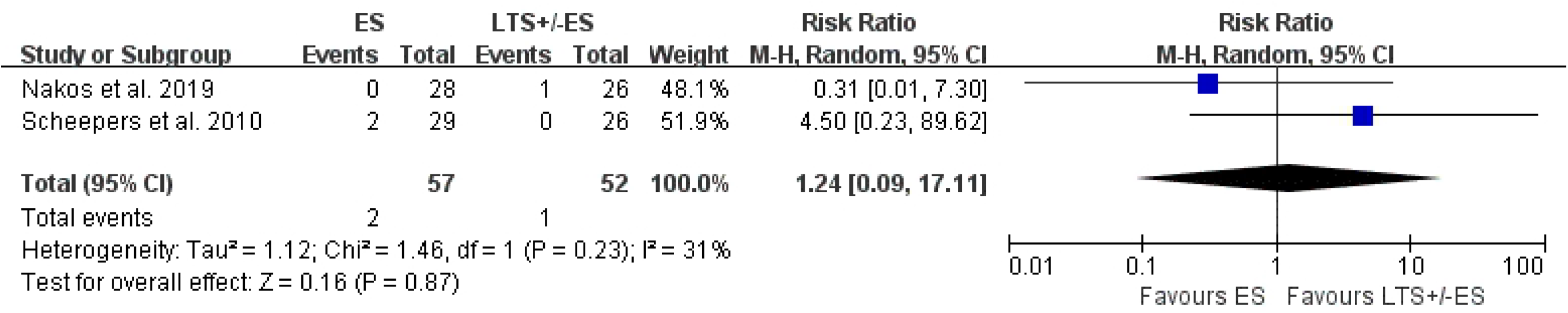
Forest plots of the effect sizes for complications at the end of the study follow-up points.

In the Nakos trial [8], one patient in the LTS group developed an abscess in the lateral canthal area at 12 months. In the Scheepers trial [9], a suture-related granuloma was observed with two patients in the ES group and no case of ectropion was observed. In the Nakos [8] and Scheepers trials [9], the risk ratios for complications between the ES group and LTS +/- ES group were 0.31 (95% CI: 0.01 to 7.30, P = 0.47) and 4.50 (95% CI: 0.23 to 89.62, P = 0.32), respectively.

A total of 10 RCTs have been performed on entropion to date. Except for the two articles included in this meta-analysis, there was one unpublished RCT (ISRCTN 29030032), three RCTs for suture materials, and four RCTs comparing surgical methods (Table 4).

**Table 4.** Characteristics of the excluded randomized controlled trials.

| **Trials** | **Country** | **Comparison arms (n)** | **Follow-up** | **Surgeon** | **Summary of results** |
| --- | --- | --- | --- | --- | --- |
| **Suture materials** |  |  |  |  |  |
| Cartmill 2014 | UK | ES using 5-0 Vicryl/7-0 Vicryl<br>(21/21) | 1 mo. | NA | Favors 7-0 Vicryl |
| Jensen 1983 | Denmark | Celsus' technique using<br>Dexon/Silk (29/20) | 1 mo. | various | Favors Dexon |
| Singh 2008 | Nepal | ES using silk/chromic catgut<br>(25/25) | 3 mo. | NA | Favors silk |
| **Surgical methods** |  |  |  |  |  |
| Dulz 2019 | Germany | QP<sup>‡</sup>/LTS+ES (36/30) | 14 mo. | single | Similar |
| Jamison 2020* | UK | LTS+Jones/LTS+ES (15/16) | 12 mo. | NA | Similar |
| Lopez-Garcia 2016 | Spain | LTS/modified LTS (46/50) | 6 yr. | single | Favors modified LTS |
| Xu 2015 | China | OMS/OMS+Skin Ex. (63/63) | 18 mo. | NA | Favors OMS+Skin Ex. |

ES, everting sutures; NA, not available; RCT, randomized controlled trial; QP, Quickert procedure; LTS, lateral tarsal strip; OMS, orbicularis oculi muscle shortening; mo., months; yr., year. \* This trial result was from a comment on Nakos 2019, <sup>‡</sup> This QP is for the correction of both vertical laxity and horizontal laxity.

Cartmill et al. [11] compared foreign body inflammatory responses in a group where 5-0 Vicryl (21 eyes) and a group where 7-0 Vicryl (21 eyes) was used to perform ES in a total of 42 eyes. The use of 7-0 Vicryl was found to significantly reduce the foreign body inflammatory response. Jensen et al. [12] compared the success rate in a group where Dexon (29 eyes) and a group where silk (21 eyes) was used when performing the Celsus technique in a total of 49 eyes and the success rate was not statistically different between the groups. Although there was no significant difference, the study reported that the immediate cosmetic satisfaction was increased and time consumption was reduced in the group sutured using Dexon, an absorbable material. Singh et al. [13] compared the success rate and the cost of surgery in groups using silk (25 eyes) or chromic catgut (25 eyes) for ES in a total of 50 eyes. Although there was no statistically significant difference in the success rate between the two groups, the use of silk significantly reduced the cost of surgery. Therefore, when selecting suture material for involutional entropion surgery, it is better to use a thinner absorbable material, but the cost of surgery should be considered in a country with low socioeconomic status.

Dulz et al. [14] performed a Quickert procedure, which corrects not only vertical laxity but also horizontal laxity, on 36 eyes, LTS and ES simultaneously (LTS + ES) on 30 eyes in a total of 66 eyes. The recurrence rates were compared and were not statistically significant between the two groups. Jamison et al. [15], in a comment on the Nakos trial [8], compared the complications and recurrence rates between a group (15 eyes) treated with LTS and the Jones procedure and a group (16 eyes) treated with LTS and ES, for a total of 31 eyes. Because of the short-term follow-up period, subtle differences were not revealed between the two groups. Lopez-Garcia et al. [16] compared the recurrence rate and the changes in horizontal laxity between a group (46 eyes) treated with LTS and a group (50 eyes) treated with modified LTS, for a total of 96 eyes. The modified LTS showed statistically significant reductions in the recurrence rate and horizontal laxity. In a total of 126 eyes, Xu [17] showed that the short-term effectiveness rate and long-term cure rates in the group (63 eyes) treated with the combination of OOM shortening and skin excision were significantly increased compared to the group (63 eyes) treated with OOM shortening alone.

## Discussion

The ES technique introduced by Quickert & Rathbun [18] in 1971 is a method for correcting vertical laxity. Several sutures are inserted through the conjunctiva, deep within the inferior fornix to evert the lower eyelid [9]. The advantage of the ES technique is simplicity. Even in patients who cannot stop anti-coagulation therapy, ES can be easily performed in outpatient clinics and can be applied to bedridden patients in conditions making it difficult to enter operating rooms. The procedure can even be performed by trained ophthalmic nurses [8]. Mohammed & Ford [19] performed nurse-led ES in 90 lids of 82 patients for involutional lower eyelid entropion at an outpatient clinic and showed a recurrence rate of 20%. The disadvantage of the ES technique is its relatively high recurrence rate. In the case of correction by ES only, the reported recurrence rates were 15% [20], 11.8% [21], and 49.3% [22].

The LTS technique introduced by Anderson & Gordy [23] in 1979 is a method for correcting horizontal laxity. This technique is performed under local anesthesia, and initially, lateral canthotomy and inferior cantholysis are performed to mobilize the lateral aspect of the eyelid. To prepare the outer tarsal strip from the lateral edge of the eyelid, the anterior lamella, that is, the skin and orbicularis muscle, and the lateral tarsus are separated, and the conjunctiva and the lid margin tissue are removed. Then, the prepared outer tarsal strip is attached to the internal lateral orbital rim. The advantage of LTS is that it has more rapid rehabilitation, better cosmetic results, and lower complications compared to other surgical techniques used to correct horizontal laxity of the lower eyelid. [24] In addition, LTS shows lower recurrence than ES. When LTS was combined with ES or Quickert’s sutures or lower eyelid retractor advancement (LERA), the recurrence rates were 18.2% [25], 12.2% [26], and 5.1% [27].

About 10 years ago, the first RCT paper [9] comparing the recurrence rates between ES and LTS + ES was published, and nine years later, an additional RCT paper comparing the ES and LTS recurrence rates was published. The meta-analysis of these two RCTs found statistically significantly lower recurrence rates for LTS +/- ES than for ES for involutional entropion. The complication rates were not significantly different from each other, and both surgical methods showed relatively low complication rates of 0 – 6.9%. Therefore, to lower the recurrence rate in patients with involutional entropion, horizontal laxity correction alone or a procedure combining LTS with or without ES, rather than correcting only vertical laxity with ES is effective.

However, Lee et al. [28] evaluated horizontal laxity. LERA was performed on patients with involutional entropion in Japan and the cases were divided into those with and without horizontal laxity. The recurrence rate of the patients with horizontal laxity was 8.7%, but there were no recurrences in patients without horizontal laxity after 22 months. In Korea, Jang et al. [22] reported that the recurrence rate was 49.3% when only ES was performed for involutional entropion with horizontal laxity. However, in China, Tsang et al. [21] reported a recurrence rate of 11.8% when only ES was performed for involutional entropion without horizontal laxity. In the RCT conducted by Scheepers et al. [9], the patients in the recurrence group had an average of 10.8 mm of horizontal laxity, more than the other involutional entropion patients, although the average horizontal laxities of the ES group (9.6 mm) and the ES+LTS group (9.5 mm) were not significantly different from each other.

That is, rather than performing LTS or LTS+ES for all involutional entropion cases, to lower the recurrence rate, horizontal laxity should be evaluated first and the appropriate surgical method should be selected accordingly. This is because there is an advantage of ES only, such as in cases where the horizontal laxity is not severe or in patients whose conditions cannot tolerate more invasive horizontal laxity correction, such as by LST. Consideration can be given to using sequential methods where ES is performed first and then when the entropion recurs, horizontal laxity correction only or a combined procedure is performed. Also, another method, such as ES for low horizontal laxity and LTS or LTS+ES for higher horizontal laxity, dependent upon the degree of horizontal laxity may be considered.

In the two RCT studies selected for this meta-analysis, there was no mention of blinding, and all were assessed as high risk of bias in quality assessment. It is more difficult to conceal group allocations in surgical intervention RCTs than medication assignments in drug RCTs, but more rigorously designed RCT studies are required to obtain more reliable results. In addition, although the characteristics of entropion are different in Asians and non-Asians [4], the two RCT studies selected for this meta-analysis were conducted only in Europe. Therefore, RCTs in Asian patients are necessary to confirm the results reported here.

## Acknowledgments

We gratefully acknowledge the help of Harrisco for providing editing services for this paper.

## S1 Appendix. PRISMA checklist

